# Structural basis for the strict substrate specificity of β-D-galactofuranosidase from *Streptomyces* sp. JHA19

**DOI:** 10.1101/2024.08.23.609336

**Authors:** Noriki Fujio, Chihaya Yamada, Toma Kashima, Emiko Matsunaga, Robert J. Nash, Kaoru Takegawa, Shinya Fushinobu

## Abstract

D-Galactofuranose (Gal*f*) is widely distributed in polysaccharides and glycoconjugates of bacteria, filamentous fungi, and protozoa. The biosynthetic and degradation pathways of Gal*f* in pathogens have attracted attention as potential targets for drug development. β-D-Galactofuranosidase (Gal*f*-ase) releases Gal*f* from the non-reducing ends of glycans. Gal*f*-ase activity is often exhibited by α-L-arabinofuranosidases, which hydrolyze a similar substrate. Several Gal*f*-specific Gal*f*-ases that cleave only Gal*f* and not L-arabinofuranose (Ara*f*) have recently been identified in the glycoside hydrolase (GH) families 2, 5, and 43. However, the structural basis of how they discriminate the substrates is unknown. ORF1110, belonging to GH2, is the first identified Gal*f*-specific Gal*f*-ase isolated from *Streptomyces* sp. JHA19. Here, we solved the crystal structure of ORF1110 in complex with a mechanism-based potent inhibitor, D-iminogalactitol (*K*_i_ = 65 μM). ORF1110 binds to the C5-C6 hydroxy groups of D-iminogalactitol with an extensive and integral hydrogen bond network. This result suggests that in the case of Ara*f*, which lacks the C6 hydroxymethyl group, this network is not formed. The domain structure of ORF1110 is similar to that of β-glucuronidases and β-galactosidases, which belong to the same GH2 family and hydrolyze pyranose substrates. However, their active site structures were completely different. A predicted structure of the C-terminal Abf domain of ORF1110 was very similar to the carbohydrate-binding module family 42, which binds Ara*f*, and pockets that may bind Gal*f* were present.

## Introduction

D-Galactofuranose (Gal*f*) is a pentacyclic sugar formed from D-galactose and distributed as glycoconjugates and polysaccharides in bacteria, filamentous fungi, and protozoa [1,2]. Gal*f*-containing glycans of pathogenic microbes and parasitic protozoa, such as *Aspergillus fumigatus* [3], *Mycobacterium* [4], *Trypanosoma* [5], and *Leishmania* [6], have been studied in detail. As Gal*f* is absent in mammals and higher plants, its metabolic pathways in pathogens have attracted interest as targets for drug development to combat microbial infections. For example, Gal*f* constitutes approximately 5% of the dry weight of the filamentous fungus *A. fumigatus* [3] and plays a significant role in pathogenicity [7–9]. Biosynthetic enzymes for Gal*f*-containing glycoconjugates, e.g., galactofuranosyltransferases, have been also extensively studied [10,11], while studies of degradative enzymes have only intensified in recent years.

β-D-Galactofuranosidase (Gal*f*-ase, EC 3.2.1.146) is an enzyme that releases Gal*f* from polysaccharides and glycoconjugates [11]. Gal*f*-ases have been found in several glycoside hydrolase (GH) families in the Carbohydrate-Active enZyme (CAZy) database [12], namely in GH2 [13], GH5 [14], GH43 [15], GH117 [16], GH159 [16], and GH182 [17]. Because Gal*f* is structurally similar to α-L-arabinofuranose (Ara*f*) (Fig. 1), some α-L-arabinofuranosidases (Ara*f*-ases, EC 3.2.1.55) also exhibit Gal*f*-ase activity, as exemplified by the bifunctional *Aspergillus niger* Ara*f*-ases in GH51 and GH54 [18]. The gene responsible for Gal*f*-specific Gal*f*-ase from *Streptomyces* sp. JHA19 (ORF1110) was first identified in 2015 [13]. ORF1110 belongs to GH2 and exhibited exclusive substrate specificity towards *p*-nitrophenyl (*p*NP)-β-D-Gal*f* with kinetic parameters of *K*_m_ = 0.25 mM and *k*_cat_ = 3.5 s^−1^ but without activity towards *p*NP-α-L-Ara*f* [19]. Subsequent studies have identified both Gal*f*-specific and bifunctional enzymes from *Streptomyces* and *Aspergillus* species [20–23].

**Fig. 1.**
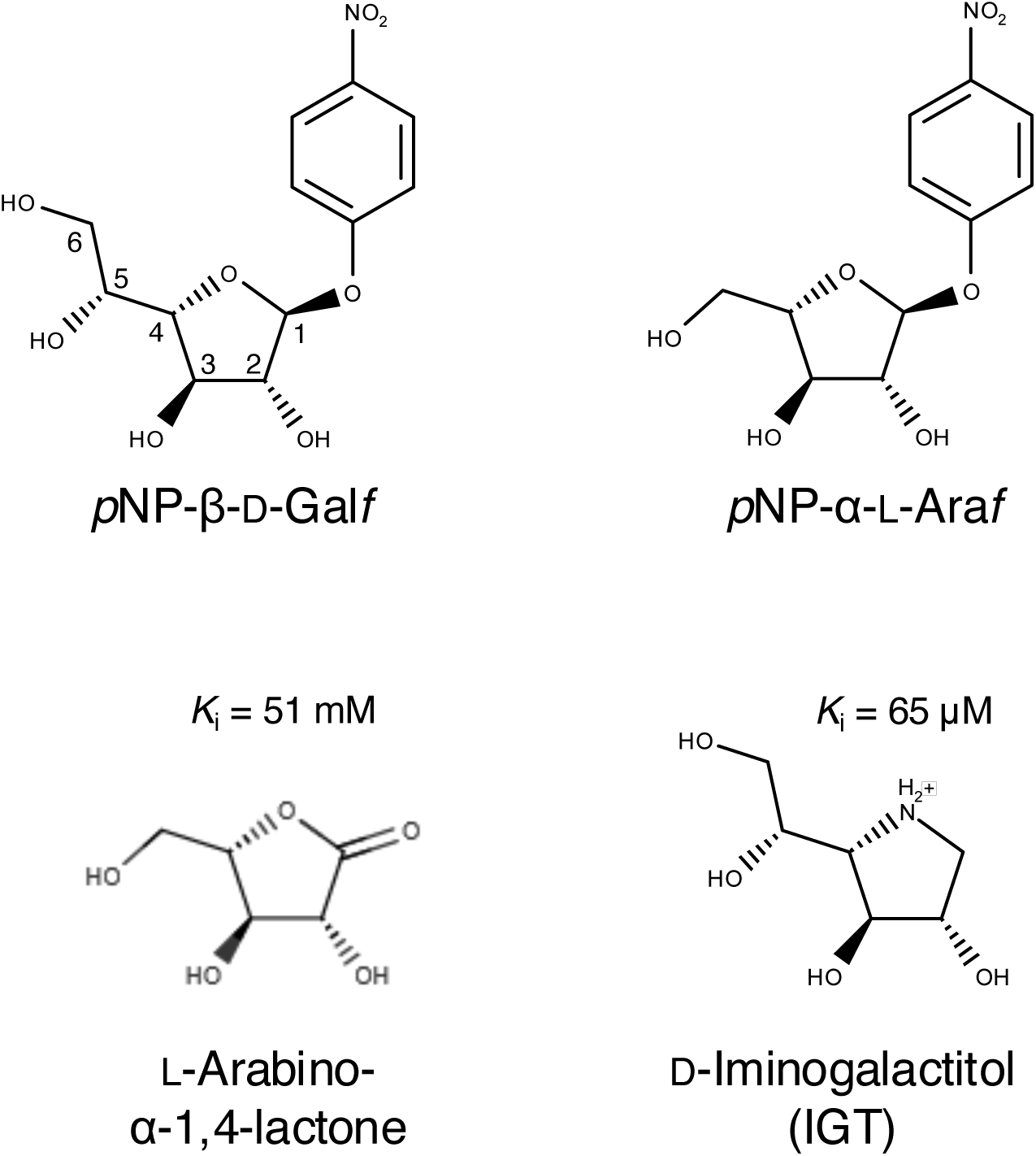
Chemical structures of *p*NP-β-D-Gal*f, p*NP-α-L-Ara*f*, L-arabino-α-1,4-lactone, and D-iminogalactitol (IGT). The calculated p*K*_a_ of the imino group of IGT is 7.24. The *K*_i_ values for ORF1110 are shown for the inhibitors.

Recently, Gal*f*-specific Gal*f*-ases were also found in GH5 subfamily 13 (GH5_13) from *Mycobacterium tuberculosis* (GlfH1, Rv3096) [14] and GH43_34 from *A. niger* (XynD) [15]. However, the three-dimensional structures of these enzymes have not yet been reported. A crystal structure of a bifunctional Ara*f*-ase/Gal*f*-ase from *Caldicellulosirputor hydrothermalis* belonging to GH159 was reported, but the structure was in a ligand-free (apo) form [24]. Therefore, no structural basis for the substrate specificity of Gal*f*-ases based on experimental complex structures with ligands is available. In this study, we determined the crystal structure of ORF1110 in complex with a deoxynojirimycin-like inhibitor of Gal*f*, revealing the structural basis for its strict substrate specificity.

## Results and Discussion

### The protein construct and an inhibitor for crystallography

ORF1110 is a 786 amino acid protein with a signal peptide of 44 amino acids at the N-terminus, according to the SignalP-6.0 server (Fig. S1A) [25]. A Pfam domain search [26] suggested that ORF1110 consists of three GH2-related domains (2N, GH2, and 2C, residues 86-203, 252-339, 382-492) and C-terminal α-L-arabinofuranosidase B (AbfB) domain (residues 641-780). The AlphaFold server [27] predicted that residues 45-635 fold as an integral unit, and AbfB is a separate domain with a β-trefoil fold. Therefore, we prepared an N-terminally histidine-tagged protein construct of residues 45-635 of ORF1110 for structural analysis.

L-Arabino-α-1,4-lactone (Fig. 1) was shown to inhibit ORF1110 but with a high *K*i value (51 mM) [13], probably due to a lack of the C6 hydroxymethyl group required for binding affinity to the Gal*f*-specific enzyme. Alkyl, thioheteroaryl, and thioimidoyl β-D-galactofuranoside compounds also inhibited ORF1110, but with moderate *K*_i_ values ranging from 0.4 to 3.0 mM [19]. In this study, we employed a 1-deoxynojirimycin-like derivative of Gal*f*, D-iminogalactitol (IGT, Fig. 1), which inhibited UDP-galactopyranose mutase but was inactive against mycobacterial UDP-Gal*f* transferases [28,29], for the crystallographic study. When assayed with *p*NP-β-D-Gal*f*, IGT competitively and potently inhibited ORF1110 with *K*_i_ = 65 μM (Fig. S2).

### Crystal structure

The crystal structure of ORF1110 (45-635) was solved by the single-wavelength anomalous dispersion method using selenomethionine (SeMet)-labeled protein, and we determined the structures of apo (glycerol complex, 1.80 Å resolution) and IGT-complexed (1.70 Å) forms (Table 1). Both structures contained two ORF1110 molecules in the asymmetric unit (Fig. S3A). There are many water molecules between the two monomers in the asymmetric unit, and no strong direct interactions were observed with the symmetry-related molecules. The PISA server [30] predicted that the biological assembly in solution is a monomer. Each molecule in the crystal structure contained one Mg^2+^ ion, which is coordinated by the side chains of D157, D201, and Q213, the main chain carbonyl of A231, and two water molecules (Fig.S3B). The CheckMyMetal server indicated that the coordination distances and geometry are typical of Mg^2+^ binding sites, but not Ca^2+^ or other metal binding sites of proteins [31].

**Table 1.** Crystallographic data statistics of Gal*f*-ase.

| Data set | Se-Met | Apo (glycerol)<br>Y585H SeMet | D-iminogalactitol<br>(IGT) |
| --- | --- | --- | --- |
| **Data collection** |  |  |  |
| Beamline | AR-NW12A | BL1A | AR-NW12A |
| Wavelength (Å) | 0.9750 | 1.052 | 1.0000 |
| Space group | $P2_1$ | $P2_1$ | $P2_1$ |
| Unit cell (Å, °) | $a = 82.466, b = 57.359,$<br>$c = 109.970, \beta = 94.766$ | $a = 84.551, b = 57.346,$<br>$c = 110.476, \beta = 92.048$ | $a = 82.364, b =$<br>$57.346, c = 109.732, \beta$<br>$= 94.934$ |
| Resolution (Å) | 47.45–2.00 (2.04–2.00) | 47.45–1.80 (1.83–1.80) | 47.41–1.70 (1.73–1.70) |
| Total reflections | 960,340 (57,876) | 340,935 (16,720) | 383,479 (18,084) |
| Unique reflections | 69,636 (4,450) | 98,339 (4,807) | 110,248 (5,375) |
| $R_{\text{merge}}$ | 0.197 (0.866) | 0.088 (0.525) | 0.097 (0.641) |
| $R_{\text{pim}}$ | 0.055 (0.248) | 0.055 (0.328) | 0.061 (0.409) |
| Mean $I/\sigma(I)$ | 12.6 (3.2) | 9.8 (2.6) | 9.0 (1.9) |
| $CC_{1/2}$ | 0.996 (0.849) | 0.991 (0.756) | 0.995 (0.658) |
| Completeness (%) | 100.0 (100.0) | 100.0 (99.9) | 98.2 (96.8) |
| Multiplicity | 13.8 (13.0) | 3.5 (3.5) | 3.5 (3.4) |
| Anomalous completeness (%) | 99.9 (100.0) |  |  |
| Anomalous multiplicity | 7.0 (6.5) |  |  |
| **Refinement** |  |  |  |
| Resolution (Å) |  | 47.49–1.80 | 47.46–1.70 |
| No. of reflections |  | 98,321 | 110,232 |
| $R_{\text{work}}/R_{\text{free}}$ | | 0.143/0.177 | 0.153/0.189 |
| Number of atoms |  |  |  |
| Amino acids |  | 8,877 | 9,012 |
| Ions |  | 2 | 2 |
| Ligands |  | 137 | 78 |
| Waters |  | 816 | 797 |
| B-factors (Å<sup>2</sup>) |  |  |  |
| Amino acids |  | 15.6 | 16.13 |
| Ions |  | 2 | 11.58 |
| Ligands |  | 137 | 26.6 |
| Waters |  | 816 | 23.45 |
| Clashscore |  | 1.52 | 2.68 |
| Molprocity score |  | 1.14 | 1.22 |
| RMSD from ideal values |  |  |  |
| Bond lengths (Å) |  | 0.0154 | 0.0155 |
| Bond angles (°) |  | 2.14 | 2.12 |
| Ramachandran plot (%) |  |  |  |
| Favored |  | 97.05 | 97.22 |
| Allowed |  | 2.69 | 2.61 |
| Outlier |  | 0.26 | 0.17 |
| PDB code |  | 9J6M | 9J6N |

Values in parentheses are for the highest resolution shell.

The overall structure of ORF1110 (Fig. 2A) consists of the N-terminal jellyroll domain (residues 45-244), the middle Ig-like domain (245-360), and the C-terminal TIM barrel domain (361-626). The active site is located at the center of the TIM barrel domain, where IGT was bound in the complexed form (Fig. 2B). The furanose-like pentacyclic ring of IGT adopts a ^4^*T*_O_ conformation (*P* = 249° and *φ*_m_ = 44.6° for chain A, and *P* = 250° and *φ*_m_ = 47.4° for chain B) as analyzed by the Altona-Sundaralingam phase angle parameter [32]. The apo crystal form contained a glycerol molecule in the active site because the crystal was cryoprotected in the presence of 20% glycerol (Fig. S3C). The glycerol molecule is hydrogen-bonded with the side chains of H401, K212, E594, and Y551. The main chain structures of both chains (A and B) in the two crystal forms were virtually the same, and the Cα root mean square deviation (RMSD) was less than 0.231 Å for all chain pairs. Therefore, we mainly describe chain A of the IGT complex.

**Fig. 2.**
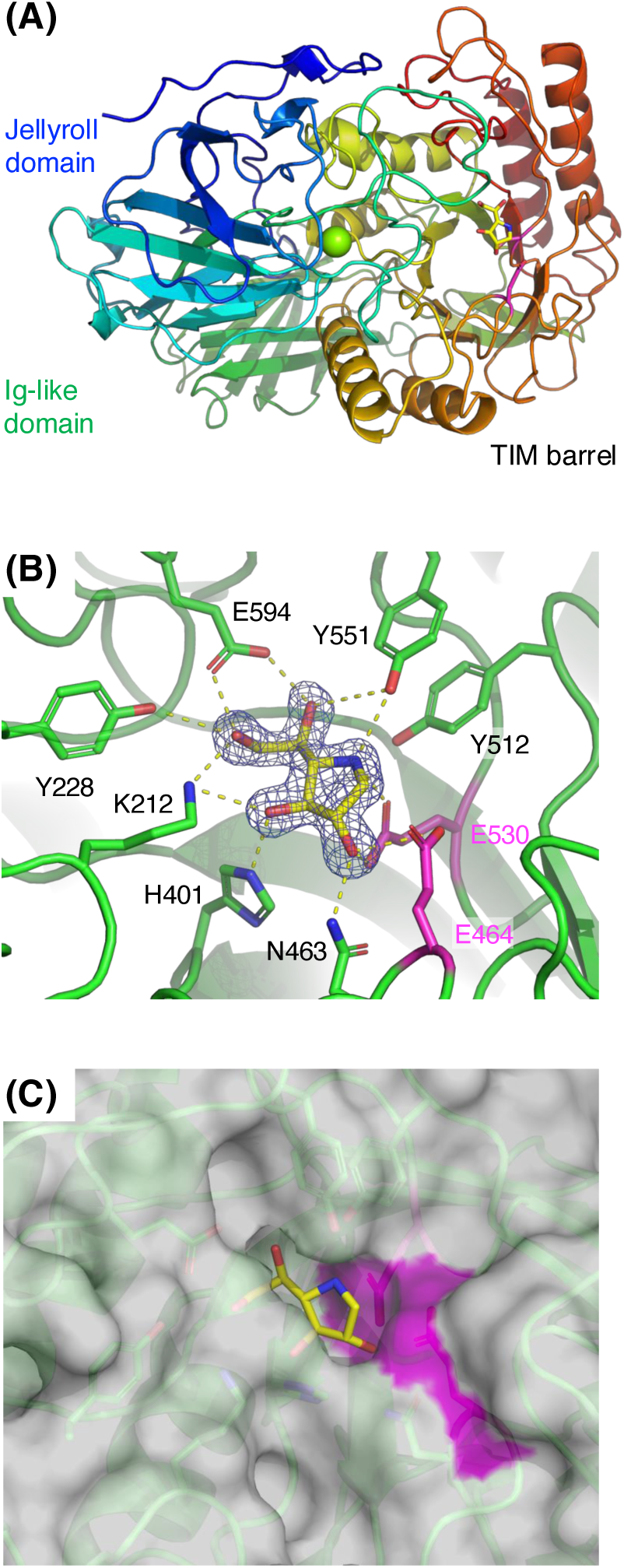
Crystal structure of ORF1110 (residues 45-635) complexed with IGT. (A) Overall structure. The polypeptide is shown in rainbow color. The catalytic residues, IGT, and Mg^2+^ ion are shown as magenta and yellow sticks and a green sphere, respectively. (B) Active site. A Polder map (4σ) of IGT is shown. Hydrogen bonds are shown as yellow dotted lines. The catalytic and other residues in the active site are shown as magenta and green sticks, respectively. (C) Molecular surface near the active site. The surface area formed by the catalytic residues is colored magenta.

The IGT molecule is recognized by the protein with direct hydrophilic interactions without mediating water molecules, and there is no stacking interaction with an aromatic residue (Fig. 2B). The catalytic residues of ORF1110 are E530 (nucleophile) and E464 (acid/base catalyst), as inferred by sequence alignment with GH2 β-galacto(pyrano)sidases (EC 3.2.1.23) [33,34]. Both E530 and E464 are located near the C1 atom of IGT and are suitably positioned for nucleophilic attack to the anomeric carbon of Gal*f* and proton donation at the scissile glycosidic bond oxygen, respectively. Mutation of E530 and E464 to alanine significantly reduced the activity in our previous study [13]. The endocyclic imino group of IGT forms a hydrogen bond with Y551. This imino group is expected to be protonated at the optimum pH (4.5) [19]. The electrostatic interaction with the negatively charged catalytic residues of GHs usually contributes to the high potency of mechanism-based iminosugar inhibitors, which mimic the partial positive charge at the oxocarbenium-like transition state [35]. The O2 hydroxy forms a hydrogen bond with N463, and the O3 hydroxy forms hydrogen bonds with H401 and K212. The most interesting structural feature of the IGT complex is the hydrogen bonding interactions of the C5-C6 hydroxy groups. The O5 hydroxy is hydrogen-bonded with Y551 and E594, and the O6 hydroxy is recognized by three residues, K212, Y228, and E594. Thus, E594 forms bidentate hydrogen bonds with the two hydroxy groups. Our previous study reported that the mutation of E594 to alanine significantly reduced its activity [13]. In the case of Ara*f*, which lacks the C6-O6 hydroxymethyl group of Gal*f*, three of the five hydrogen bonds are lost with the O5-O6 exocyclic group, and the integrity of the active site interaction will be significantly impaired. ORF1110 has an open active site that is typically present in exo-glycosidases, and there are no positive subsites (Fig. 2C). This molecular feature is consistent with the enzyme’s ability to hydrolyze *A. fumigatus* galactomannan [13], which has β-1,3, β-1,6, and β-1,5-linked Gal*f* oligosaccharide side chains on its galactan main chain [2].

### Comparison with GH2 enzymes

The CAZy database currently lists 16 enzymatic activities, including β-galactosidase, β-glucuronidase (GUS, EC 3.2.1.31), β-galacturonidase (EC 3.2.1.67), and Gal*f*-ase in the GH2 family (http://www.cazy.org/GH2.html). A Dali structural similarity search revealed that ORF1110 mostly resembles the uncharacterized BACCELL_01794 protein from *Bacteroides cellulosilyticus* (Table S1). The second hit was GH2 GUS from *Bacteroides thetaiotaomicron* (BtGUS) [36], which has 43% sequence identity with ORF1110. Subsequent hits were bacterial GH2 GUSs with around 20% sequence identity. GH2 β-galactosidases have lower structural similarity to ORF1110. GUSs have three-domain structures (jellyroll, Ig-like, and TIM barrel domains) similar to ORF1110, whereas β-galactosidase II from *Bacillus circulans* (β-Gal-II) [37] has two additional domains (DUF4982 and bacterial Ig-like I) at the C-terminus (Fig. S4).

The active site of ORF1110 Gal*f*-ase was compared with that of GUS from *Ruminococcus gnavus* (RgGUS) complexed with β-D-glucuronate [38] and β-Gal-II complexed with β-1,4-linked galactobiose [37] (Fig. 3). The two catalytic glutamate residues (E418 and E512 in RgGUS), one asparagine (N417 in RgGUS), and one histidine (H335 in RgGUS) are conserved in the three GH2 enzymes, but other residues in the active site are not. Most strikingly, while an aromatic stacking residue plays an important role in substrate binding at subsite –1 in GUS and β-galactosidase (W557 in RgGUS and W550 in β-Gal-II), no corresponding residue is present in ORF1110. This structural feature indicates that the GH2 family, which was established in the earliest years of CAZy classification [39] and counts more than 48,000 members in the database, acquired widely diverged active site architectures during molecular evolution from its ancestral protein.

**Fig. 3.**
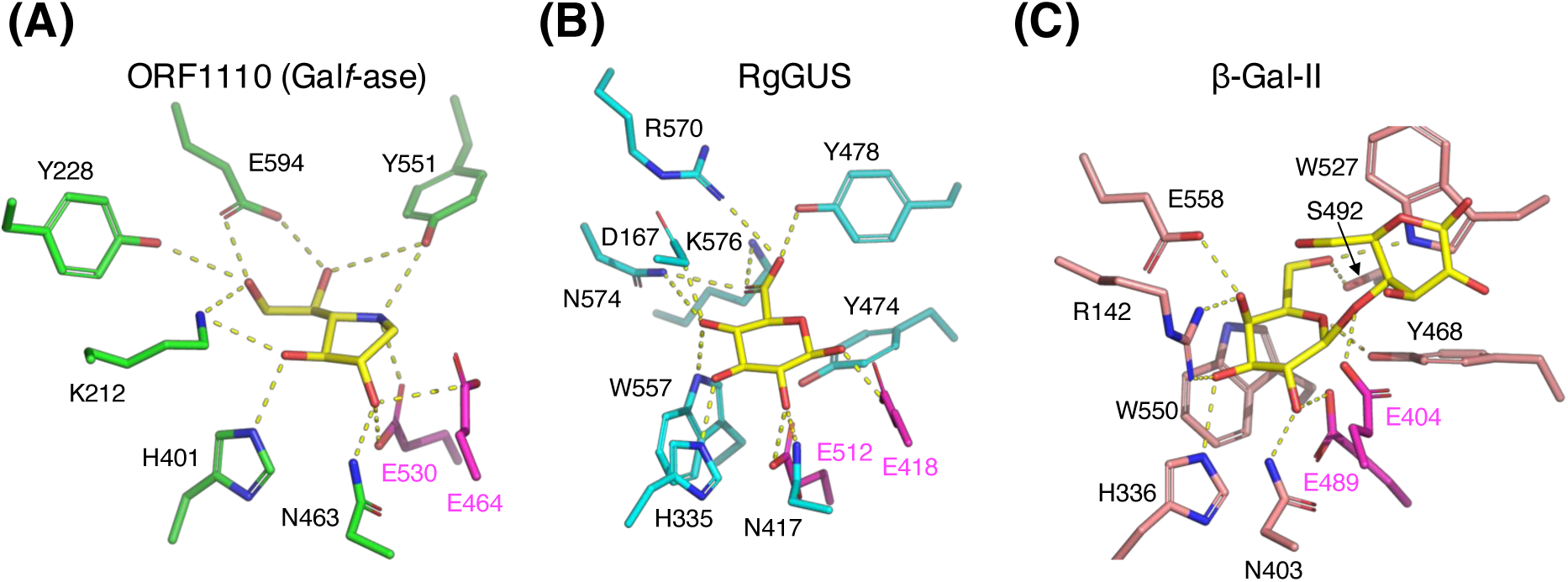
Structural comparison of the active site with GH2 β-glucuronidase (GUS) and β-galactosidase (β-Gal). (A) ORF1110 (green) complexed with IGT. (B) GUS from *Ruminococcus gnavus* (RgGUS, cyan) complexed with β-D-glucuronate (PDB ID: 6JZ5). (C) β-Gal-II from *Bacillus circulans* (pink) complexed with Gal-β1,4-Gal (PDB ID: 7CWD). The ligands and catalytic residues are shown as yellow and magenta sticks, respectively.

### Putative sugar binding sites in the AbfB domain

The C-terminal AbfB domain (residues 45-635) with a β-trefoil fold was annotated as Carbohydrate-Binding Module (CBM) family 42 by the dbCAN3 server [40]. CBM42 was originally identified as an L-arabinofuranose-binding domain (ABD) in GH54 Ara*f*-ase from *Aspergillus kawachii* (AkAbfB) [41]. Fig. 4A shows a superimposition of the ABD in AkAbfB (rainbow color) and the predicted structure of ORF110 (AbfB domain in white and TIM barrel domain in orange). These structures superimposed well with RMSD = 0.623 Å for 129 Cα atoms. Interestingly, the molecular surface of the predicted structure of the full-length ORF1110 protein exhibited two concave pockets in the AbfB domain corresponding to the two arabinose binding sites of ABD in AkAbfB (Fig. 4B). This molecular feature suggests that the AbfB domain of ORF1110 has Gal*f*-binding sites. Further biochemical and structural analyses are required to elucidate the sugar-binding function and specificity of AbfB.

**Fig. 4.**
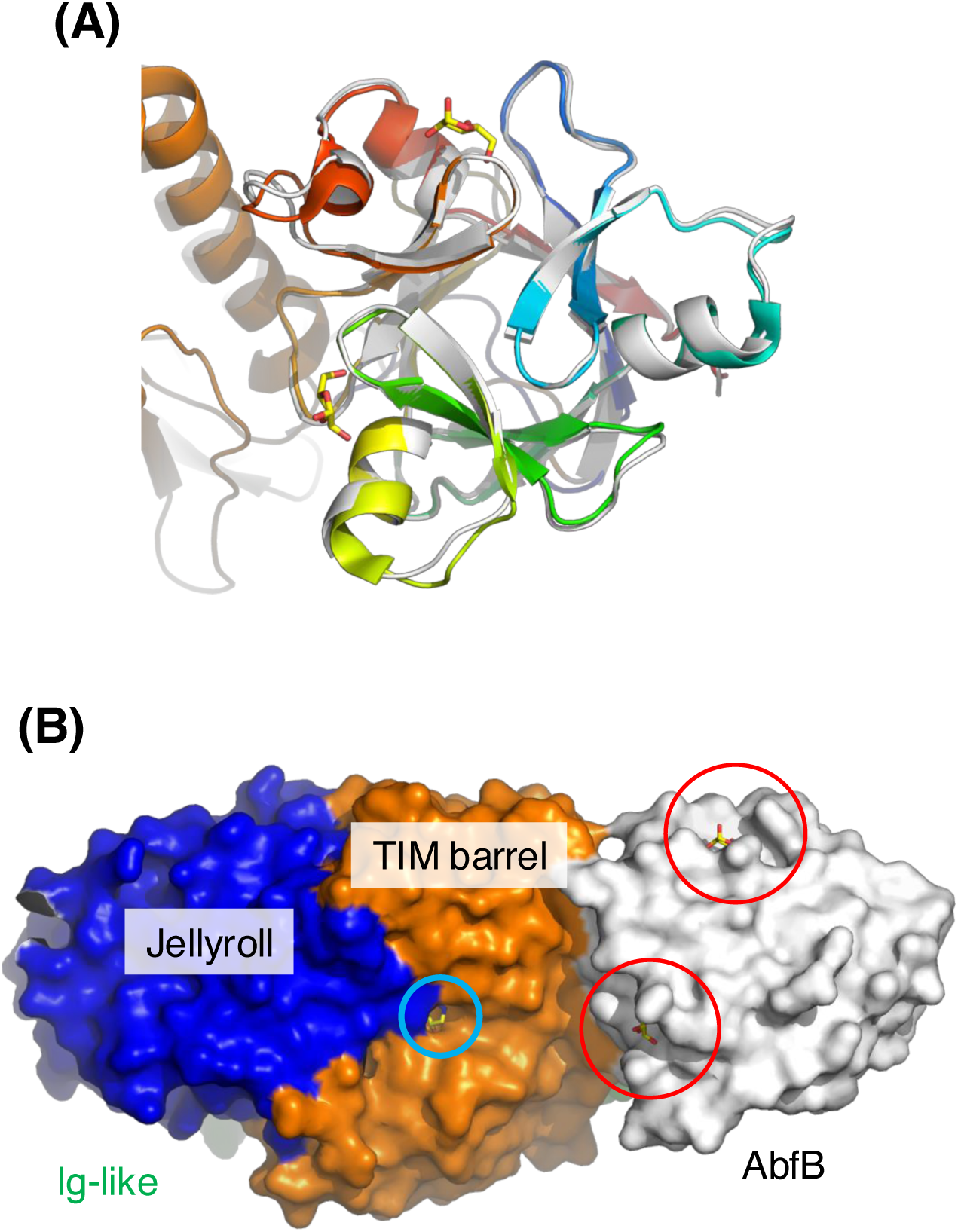
Putative sugar binding sites in the AbfB domain of ORF1110. (A) Superimposition of the predicted structure of ORF1110 (AbfB domain in white and TIM barrel domain in orange) and the L-arabinfuranose-binding domain of GH54 Ara*f*-ase from *Aspergillus kawachii* (AkAbfB) (rainbow color). L-Arabinose molecules bound to AkAbfB are shown as yellow sticks. (B) Molecular surface of the predicted structure of ORF1110 without the N-terminal signal sequence. The jellyroll (residues 45-244, blue), Ig-like (245-360, green), TIM-barrel (361-635, orange), and AbfB (636-786, white) domains are differently colored. IGT bound to the crystal structure of ORF1110 and L-arabinose molecules bound to AkAbfB are shown as yellow sticks. The red and cyan circles indicate the putative Gal*f*-binding and catalytic sites, respectively.

## Conclusion

In this study, we revealed the structural basis for Gal*f*-specific Gal*f*-ase by the determination of a complex structure with a mechanism-based potent inhibitor. It is intriguing whether other Gal*f*-specific Gal*f*-ases in GH5_13 and GH43_34 also have a similar structural system to discriminate between Gal*f* and Ara*f*. ORF1110 has an open active site that can accommodate side-chain Gal*f* units in polymeric substrates such as fungal galactomannan. β-Linked Gal*f* units were also found at the non-reducing termini of *N*-and *O*-glycans of glycoproteins, glycosphingolipids, and glycoinositolphospholipids in fungi and protozoans [2]. We also found potential Gal*f*-binding sites in the C-terminal CBM42 domain of ORF1110. Homologs of the GH2 Gal*f*-ases have been found in a wide range of organisms, from gram-positive bacteria to filamentous fungi, including Actinomycetota, Bacillota (previously known as firmicutes), and Ascomycota [13]. GH5_13 enzymes, including a Gal*f*-ase active on mycobacterial arabinogalactan, are exclusively found in bacteria [14]. On the other hand, GH43_34 enzymes include Gal*f*-ases and Ara*f*-ases from bacteria and fungi associated with CBM6, CBM13, CBM32, CBM42, and CBM66 domains [15]. Considering the broad distribution of Gal*f*-ases, structural analysis of these enzymes needs to be further advanced to establish novel pharmaceutical therapies against fungal pathogens such as *A. fumigatus*.

## Materials and methods

### Protein expression and purification

The N-terminally His_6_-tagged construct of ORF1110 (residues 45-635) was obtained by deleting the Nus-tag sequence from the pET50b-based expression plasmid constructed in our previous study [13]. The following primers were used for PCR: 5′-TCACCAT/TCCGCGGCTCTTGAAGTC-3′ (forward) and 5′-GCCGCGGA/ATGGTGATGGTGGTGATG-3′ (reverse), where the “/” mark indicates the position flanking the deleted Nus-tag sequence. PCR was performed using the KOD One PCR Master Mix (TOYOBO Co., Ltd., Osaka, Japan). The expression construct begins with the following amino acid sequence: MGSSHHHHHHSAA<u>LEVLFQGP</u>, where the HRV 3C protease site is underlined. Because an unintended Y585H substitutional mutation was introduced during PCR, we reverted the mutation by site-directed mutagenesis using KOD One DNA polymerase and methylation-sensitive restriction enzyme DpnI (TaKaRa Bio Inc., Kusatsu, Japan). The following primers were used for the site-directed mutagenesis of H585Y: 5′-CCTCGGTC<u>TAC</u>ACCGAGATCACGGACG-3′ (forward) and 5′-CGGT<u>GTA</u>GACCGAGGCGGACAGGCC-3′ (reverse), where the mutated codon is underlined.

SeMet-labeled and native proteins were expressed in *E. coli* BL21-CodonPlus(DE3)-RIL-X and BL21-CodonPlus (DE3)-RIL (Agilent Technologies, Santa Clara, CA, USA), respectively. The transformants were cultured in lysogeny broth (native protein) or LeMaster medium (SeMet-labeled protein) containing 10 mg/L ampicillin and 34 mg/L chloramphenicol at 37°C for 2.5-8 hours. Isopropyl-β-D-thiogalactopyranoside was added at a final concentration of 1 mM to induce protein expression. Following additional incubation at 18°C for 22-68 hours, the cells were harvested by centrifugation and suspended in 50 mM HEPES-NaOH (pH 8.0) and 300 mM NaCl. Cell extracts were obtained by sonication, followed by centrifugation to remove cell debris. The cell-free extract was first purified by nickel affinity chromatography using a Ni-NTA column (Qiagen, Hilden, Germany), and the enzyme was eluted with a stepwise increase in imidazole concentration (5 mM to 250 mM) in 50 mM HEPES-NaOH (pH 8.0) and 300 mM NaCl. The enzyme was further purified by gel filtration column chromatography using a HiLoad 16/60 Superdex 200 prep grade column (Cytiva, Marlborough, MA, USA) equilibrated with 20 mM HEPES-NaOH (pH 8.0) and 150 mM NaCl. Protein concentration was determined spectrophotometrically at 280 nm using a theoretical extinction coefficient based on the amino acid sequence.

### Crystallography

The crystals of ORF1110 were obtained at 20°C using the sitting-drop vapor diffusion method by mixing 0.6:0.4:0.2 μL of protein:reservoir:microseed solutions. First, SeMet-labeled crystals were obtained for initial phase determination. This protein had a catalytically irrelevant Y585H mutation. The protein and microseed solutions contained 2.0 mg/mL ORF1110 (SeMet-labeled Y585H) in 20 mM HEPES-NaOH (pH 8.0) and ×1000-fold diluted crushed crystal solution, respectively. The reservoir solution contained 50 mM potassium thiocyanate (KSCN) and 25% (w/v) PEG3350. The crystals were cryoprotected using a reservoir solution supplemented with 20% (v/v) PEG400. The same protein and microseed solutions were used for the apo (glycerol complex) crystals, and the reservoir solution contained 100 mM KSCN and 25% (w/v) PEG3350. The crystals were cryoprotected using a reservoir solution supplemented with 20% (v/v) glycerol. Crystals of the native (non-labeled) wild-type protein with the H585Y reverse mutation were prepared for soaking in a solution containing IGT. IGT was synthesized as reported previously [28,29]. The protein and microseed solutions contained 6.0 mg/mL ORF1110 in 20 mM HEPES-NaOH (pH 8.0) and 150 mM NaCl, and ×10000-fold diluted crushed crystal solution, respectively. The reservoir solution contained 75 mM KSCN and 25% (w/v) PEG3350. The crystals were soaked in a reservoir solution supplemented with 10 mM IGT and then cryoprotected using a solution further supplemented with 20% (v/v) 2-methyl-2,4-pentanediol.

For data collection, the crystals were flash-cooled by dipping them in liquid nitrogen. X-ray diffraction data were collected at 100 K on the beamlines at the Photon Factory of the High Energy Accelerator Research Organization (KEK, Tsukuba, Japan). Preliminary diffraction data were collected at SPring-8 (Hyogo, Japan). The datasets were processed using XDS [42] and Aimless [43]. Phase determination and automated model building were performed for the data of SeMet-labeled crystal using the CRANK2 pipeline [44]. Automated model building was achieved using BUCCANEER [45]. Manual model rebuilding and refinement were performed using Coot [46] and Refmac5 [47]. Polder maps were prepared using PHENIX [48]. Molecular graphic images were prepared using PyMOL (Schrödinger LLC, New York, NY, USA).

### Enzyme assay for determination of the inhibition constant

The enzyme reaction solution (90 μL) consisted of 45 μL enzyme, 36 μL substrate, and 9 μL inhibitor solution. All solutions contained 50 mM Na-acetate buffer (pH 4.5). The enzyme solution contained 3.38 μg/mL ORF1110 protein (residues 45-635 with no mutation) and 0.2% bovine serum albumin (TaKaRa Bio). At final concentrations, 1, 2, and 4 mM *p*NP-β-D-Gal*f* substrate, and 0, 0.05, 0.1, 0.2, and 0.4 mM IGT inhibitor were used. A mixture of the substrate and inhibitor solutions and the enzyme solution were separately preincubated at 37°C, and the reaction was started by mixing the two solutions. After incubation at 37°C for 2, 4, and 6 min, a 25 μL aliquot was sampled, and an equal volume of 1 M Na_2_CO_3_ was added to stop the reaction. The absorbance at 405 nm was measured using a microplate reader (BioTek Synergy H1). The absorbance was converted to the *p*NP concentration using a standard curve. Kinetic analysis was performed using the Enzyme-Kinetic Calculator [49].

## Abbreviations

Gal*f*: D-Galactofuranose
Gal*f*-ase: β-D-Galactofuranosidase
GH: glycoside hydrolase
CAZy: Carbohydrate-Active enZyme
Ara*f*-ase: α-L-arabinofuranosidase
*p*NP: *p*-nitrophenyl
IGT: D-iminogalactitol
SeMet: selenomethionine
RMSD: root mean square deviation
CBM: Carbohydrate-Binding Module

## Acknowledgments

We thank Prof. George W. J. Fleet (University of Oxford), Dr. Richard E. Lee (St. Jude Children’s Research Hospital), and Prof. Neil R. Thomas (University of Nottingham) for assistance in obtaining the inhibitor compound. We thank Dr. Takatoshi Arakawa, Mr. Jin Sekiguchi, Dr. Yujiro Higuchi, Dr. Matthew Barcus, and Mr. Masanari Tanuma for their valuable discussion and experimental support. We thank the staff of KEK-PF and SPring-8 for the X-ray data collection. This research was supported in part by JSPS-KAKENHI (17H03966, 21K19090, 23K23511 to KT, 21K15025 to CY, and 19H00929 and 23H00322 to SF) and the Research Support Project for Life Science and Drug Discovery (Basis for Supporting Innovative Drug Discovery and Life Science Research (BINDS)) from AMED under grant number JP22ama121001.

## Author contributions

KT and SF conceived and supervised the study; CY, TK, YH, KT, and SF designed experiments; NF, CY, and EM performed experiments; NF, CY, TK, and SF performed protein crystallography. RJN provided the material (inhibitor compound). SF wrote the manuscript; all authors reviewed the manuscript.

## Data availability statement

Atomic coordinates and structure factors of the crystal structures have been deposited in the Protein Data Bank under accession numbers 9J6M and 9J6N. Source data are provided with this paper.

