## Supplementary Tables and Figures for "Structural basis for the strict substrate specificity of β-D-galactofuranosidase from *Streptomyces* sp. JHA19"

**Table S1.** Result of DALI structural similarity search.

| Protein | Organism | Activity [a] | PDB ID (chain) | Z score | RMSD (Å) | LALI [b] | %ID [c] | Reference |
|---|---|---|---|---|---|---|---|---|
| BACCELL_ 01794 | *Bacteroides cellulosilyticus* | Not Tested | 7sf2 (B) | 53.0 | 1.3 | 565 | 43 | – |
| BtGUS | *Bacteroides thetaiotaomicron* | GUS | 7xyr (A) | 41.9 | 2.1 | 513 | 43 | [1] |
| DtGlcA | *Dictyoglomus thermophilum* | GUS | 6xxw (A) | 32.8 | 2.5 | 484 | 18 | [2] |
| RgGUS | *Ruminococcus gnavus* | GUS | 5z18 (E) | 32.5 | 2.5 | 485 | 19 | [3] |
| EtGalAse | *Eisenbergiella tayi* | GUS/GalAse | 6ncw (C) | 32.4 | 2.9 | 482 | 21 | [4] |
| Fp2GUS | *Faecalibacterium prausnitzii* | GUS | 6mvf (C) | 32.2 | 2.7 | 483 | 23 | [5] |
| BuGUS-1 – | *Bacteroides uniformis* | GUS | 6d89 (D) | 32.0 | 2.6 | 484 | 22 | [6] |
| β-Gal-II | *Bacillus circulans* | β-Gal | 7cwi (A) | 27.9 | 3.1 | 477 | 21 | [7] |

Analyzed using the DALI server (http://ekhidna2.biocenter.helsinki.fi/dali/) [8].

[a] GUS, β-glucuronidase (EC 3.2.1.31); GalAse, β-galacturonidase (EC 3.2.1.67); β-Gal, β-galactosidase (EC 3.2.1.23). [b] Number of aligned residues. [c] Sequence identity.

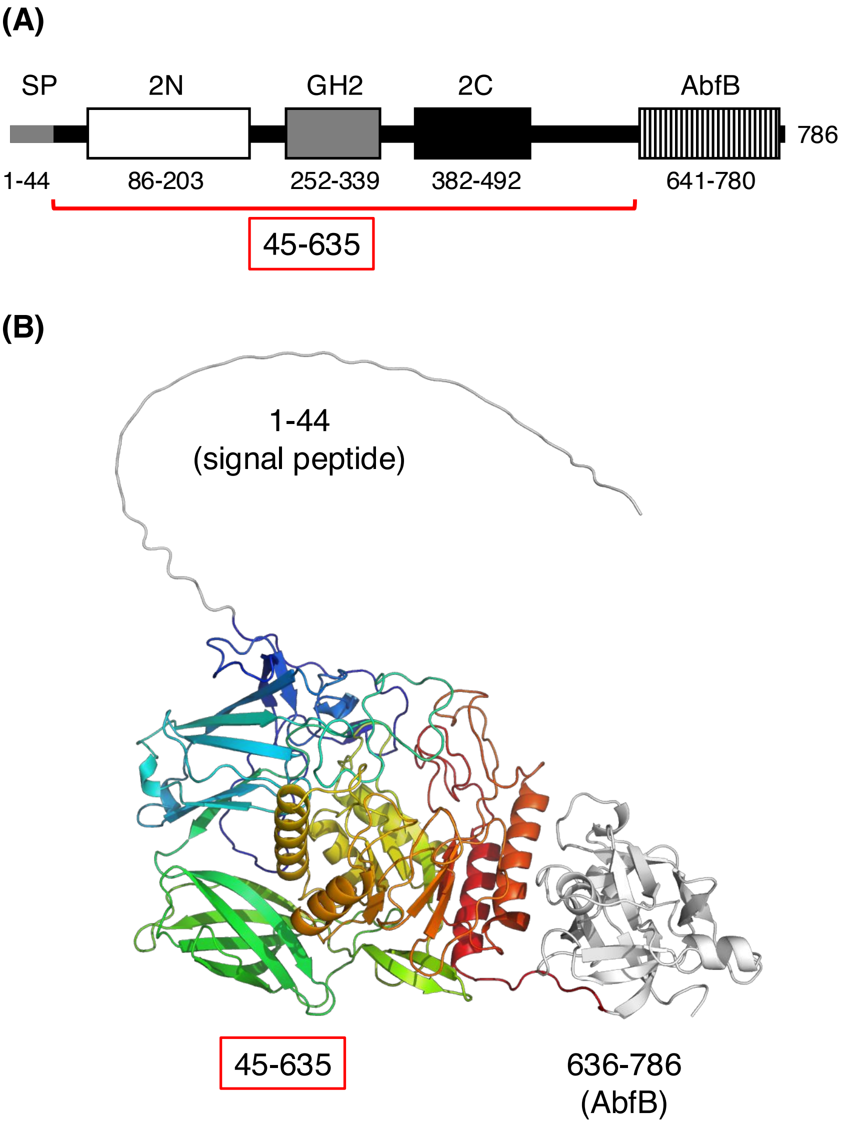


**Fig. S1.** Domain architecture and predicted structure of the full-length ORF1110 protein. (A) Domain boundaries were analyzed using Pfam [9] on the InterPro server. A red line indicates the region used for crystallization. SP, signal peptide; 2N, Glycosyl hydrolases family 2, sugar binding domain (PF02837); GH2, Glycosyl hydrolases family 2 domain (PF00703); 2C, Glycosyl hydrolases family 2, TIM barrel domain (PF02836); AbfB, α-L-arabinofuranosidase B (ABFB) domain (PF05270). The signal sequence was predicted to be residues 1 – 44 by the SignalP-6.0 server [10]. (B) A protein structure (model_0) predicted by the AlphaFold server (beta) (https://alphafoldserver.com) [11] on 7 August, 2024. The region used for crystallization is shown by rainbow color.

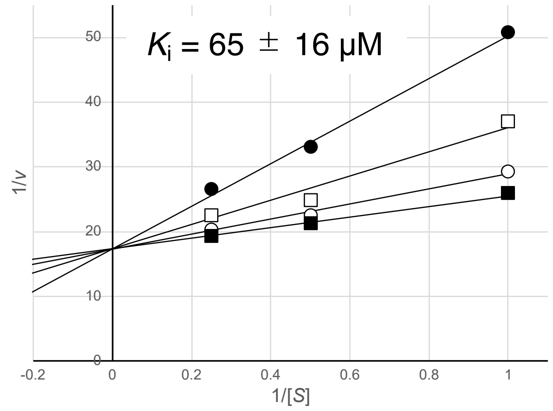


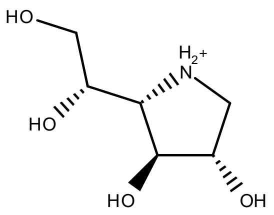


D-Iminogalactitol
(IGT)

**Fig. S2.** Double reciprocal plot showing competitive inhibition of ORF1110 by D-iminogalactitol (IGT). The hydrolytic activity of the substrate (1, 2, and 4 mM *p*NP-β-D-Gal*f*) was measured in the presence of the inhibitor (0.05, 0.1, 0.2, and 0.4 mM IGT for filled squares, open circles, open squares, and filled circles, respectively) in 50 mM Na-acetate (pH 4.5) buffer at 37°C. Kinetic analysis was performed using the Enzyme-Kinetic Calculator [12].

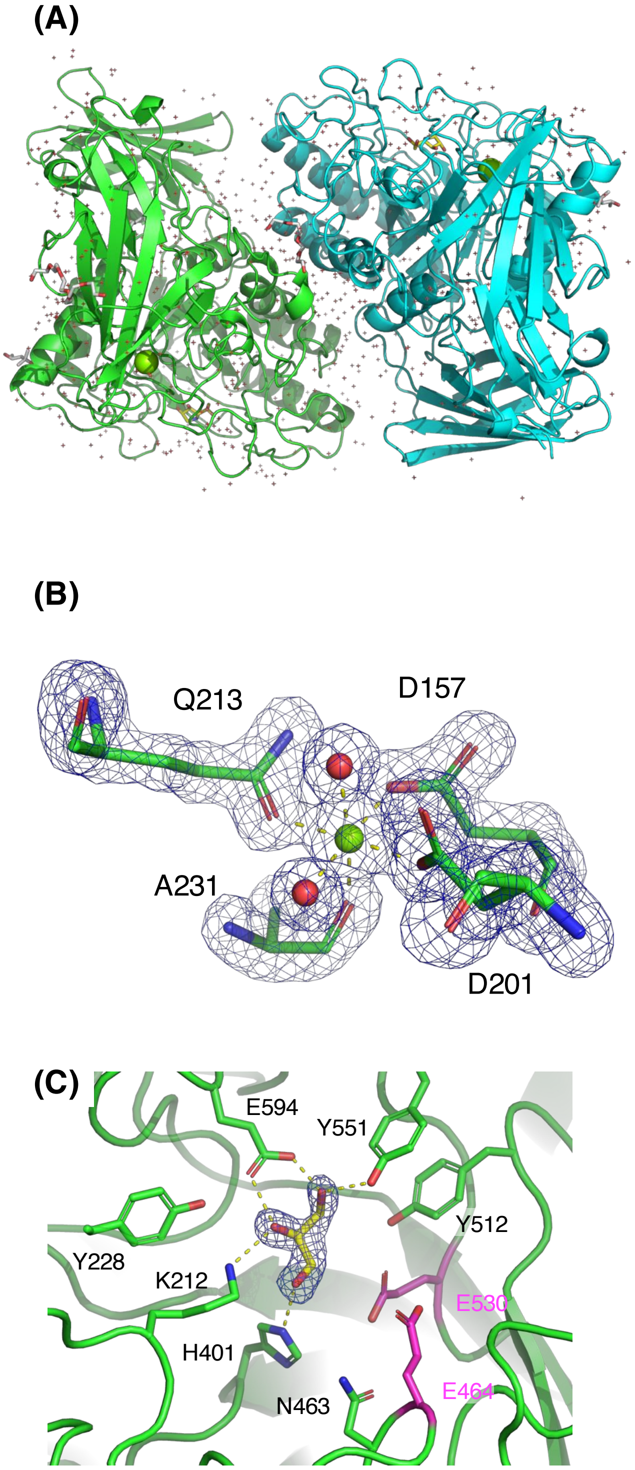


**Fig. S3.** Description of the crystal structures. (A) Dimer in the asymmetric unit (IGT complex structure). Chains A and B are shown as green and cyan cartoon models, respectively. The molecules of inhibitor (IGT), precipitant and cryoprotectant bound on the protein surface (tetraethylene glycol and hexaethylene glycol derived from polyethylene glycol reagent and 2-methyl-2,4-pentanediol), $Mg^{2+}$ ion, and water molecules are shown as yellow sticks, grey sticks, green spheres, and red crosshairs, respectively. (B) $Mg^{2+}$ binding site. A polder map of the four metal-coordinating residues (green sticks) and two water molecules (red spheres) is shown (4σ). The site on the A chain of the IGT complex is shown here although this site is present in all ORF1110 molecules in the crystal structures. (C) A Polder map (4σ) of a glycerol molecule bound to the active site is shown. The catalytic residues and other active site residues are shown as magenta and green sticks, respectively. Hydrogen bonds are shown as yellow dotted lines.

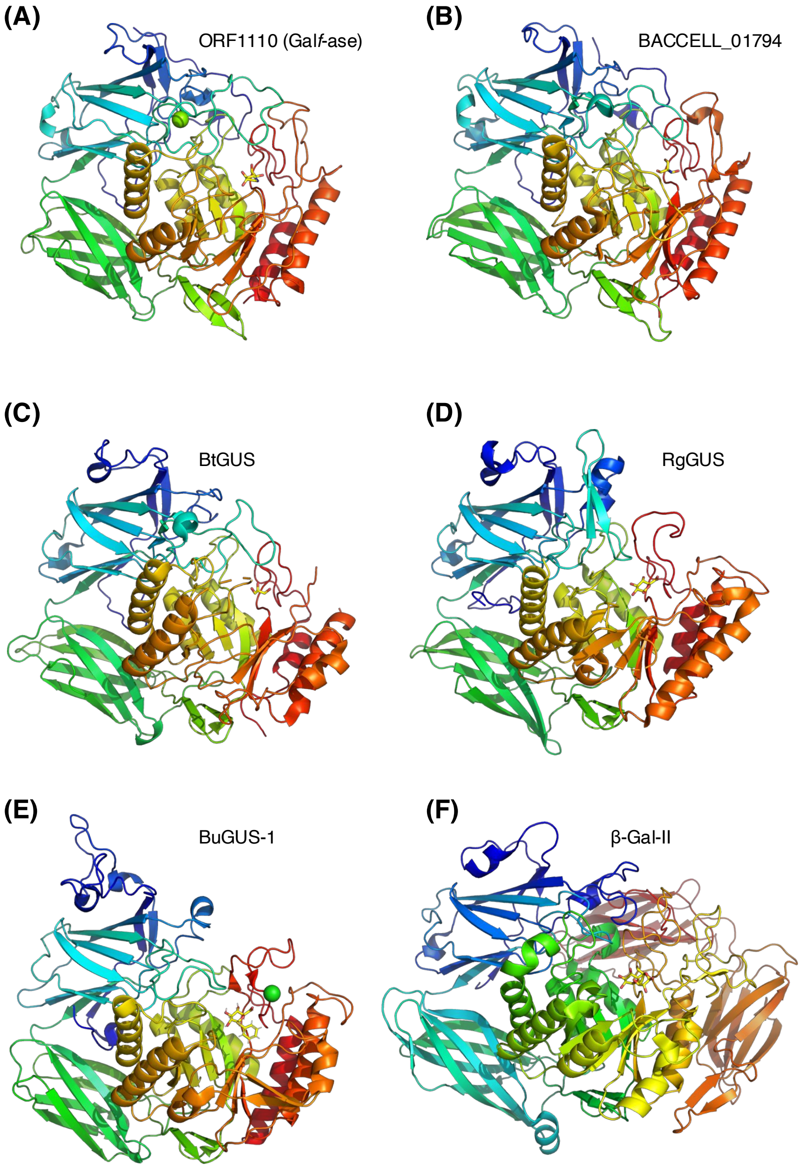


**Fig. S4.** Structural comparison of Gal*f*-ase, β-glucuronidases (GUS), and a β-galactosidase belonging to GH2. (A) ORF1110 complexed with IGT. (B) Putative protein BACCELL_01794 from *B. cellulosilyticus* complexed with Tris (PDB ID: 7SF2). (C) GUS from *B. thetaiotaomicron* (BtGUS) complexed with glycerol (PDB ID: 7XYR). (D) GUS from *R. gnavus* (RgGUS) complexed with β-D-glucuronate (PDB ID: 6JZ5). (E) GUS from *Bacteroides uniformis* (BuGUS-1) complexed with thiophenyl-β-D-glucuronide (PDB ID: 6D7F). (F) β-Galactosidase from *B. circulans* (β-Gal-II) complexed with Gal-β1,4-Gal (PDB ID: 7CWD).
